# Machine learning and mechanistic modeling for prediction of metastatic relapse in early-stage breast cancer

**DOI:** 10.1101/634428

**Authors:** C. Nicolò, C. Périer, M. Prague, C. Bellera, G. MacGrogan, O. Saut, S. Benzekry

**Author notes:** **Corresponding author:** S. Benzekry, Inria Bordeaux Sud-Ouest, 200, av. de la Vieille Tour, 33405 Talence. **Funding:** This study was achieved within the context of the Laboratory of Excellence TRAIL ANR-10-LABX-57.

## Abstract

**Purpose:** For patients with early-stage breast cancer, prediction of the risk of metastatic relapse is of crucial importance. Existing predictive models rely on agnostic survival analysis statistical tools (e.g. Cox regression). Here we define and evaluate the predictive ability of a mechanistic model for the time to metastatic relapse.

**Methods:** The data consisted of 642 patients with 21 clinicopathological variables. A mechanistic model was developed on the basis of two intrinsic mechanisms of metastatic progression: growth (parameter *α*) and dissemination (parameter *μ*). Population statistical distributions of the parameters were inferred using mixed-effects modeling. A random survival forest analysis was used to select a minimal set of 5 covariates with best predictive power. These were further considered to individually predict the model parameters, by using a backward selection approach. Predictive performances were compared to classical Cox regression and machine learning algorithms.

**Results:** The mechanistic model was able to accurately fit the data. Covariate analysis revealed statistically significant association of *Ki67* expression with *α* (p=0.001) and *EGFR* with *μ* (p=0.009). Achieving a c-index of 0.65 (0.60-0.71), the model had similar predictive performance as the random survival forest (c-index 0.66-0.69) and Cox regression (c-index 0.62 - 0.67), as well as machine learning classification algorithms.

**Conclusion:** By providing informative estimates of the invisible metastatic burden at the time of diagnosis and forward simulations of metastatic growth, the proposed model could be used as a personalized prediction tool of help for routine management of breast cancer patients.

## Introduction

Breast cancer is the most frequent and second leading cause of cancer death in women^1^. In the majority of cases, the disease is diagnosed at the early stage, when all detectable lesions, confined to the breast or nearby lymph nodes, can be surgically removed^2^. However, approximately 20-30% of patients are reported to relapse with distant metastases after surgery^3, 4^, suggesting that clinically occult micro-metastases might already be present at the time of surgery. Accurate prediction of the risk of metastatic relapse is critical to personalize adjuvant treatment and avoid use of toxic and costly therapies when not needed.

In the era of artificial intelligence, prognostic models are playing an increasing role for such a task^5^. Online tools, such as the Adjuvant!^6, 7^ and PREDICT models^8^, compute individualized survival probabilities based on multivariate statistical analysis and integration of clinical variables (age, tumor size, histological grade, hormone receptor status and nodal involvement)^5^. These tools, however, are based on agnostic statistical models, such as Cox regression^8, 9^. More recently, machine learning algorithms have started to be used^10^. Although traditionally designed for classification or regression tasks, adaptations to survival analysis include elastic net for Cox models^11^ or the random survival forests algorithm^12^. Deep learning has also recently been proposed for survival prediction from genomic data sets^13^. Nevertheless, few studies have so far investigated machine learning for prediction of breast cancer survival or recurrence^13–15^.

Mechanistic modeling approaches - where biological knowledge is used to build a simulation model - have been developed to describe metastatic dynamics^16–20^. However, none of these mechanistic models has yet been implemented as a personalized predictive tool of metastatic relapse^5^. In previous work, we evaluated a mechanistic model of metastatic development^21^ using experimental data from orthosurgical mouse models of breast cancer^22^. The model was able to describe longitudinal growth of the total metastatic burden. For human data, the model could fit size-dependent probability of 20-years metastatic relapse in a historical dataset of 2,648 breast cancer patients^22, 23^.

In the current work, we build on our descriptive model to propose an actionable tool for individualized predictions of the time to metastatic relapse (TTR), as well as reconstruction of the past natural history and prediction of future evolution of the disease. To train and validate the model, we relied on a dataset containing TTR and 21 clinical/pathological characteristics for 642 early-stage breast cancer patients. We first show a random survival forest analysis^12^, which allowed us to select a restricted number of predictors of interest. We then present the main novelty of this work: the calibration – using mixed-effects learning^24^ – of the mechanistic model. We illustrate the possible value of the mechanistic approach by performing predictive simulations of the entire cancer history of real patients, calibrated from data available at diagnosis only. Finally, we compare our results with predictive performances of classification machine learning algorithms for 5-years metastatic relapse.

## Methods

### Description of the data

The consisted of data of 642 women diagnosed with primary operable invasive breast carcinoma treated at the Bordeaux Bergonié institute between 1989 and 1993. This dataset has been comprehensively analyzed using standard statistical tools (Cox regression)^25^. Patients in this analysis did not receive any adjuvant hormone or chemotherapy. The time to metastatic relapse (TTR) was defined as the time from the date of diagnosis to the date of distant relapse. Patients with no metastasis were censored at the date of last news or death^26^. Clinical/pathological variables available in the dataset included age at diagnosis, menopausal status, histological grade, pathological T (tumor size) and N (axillary lymph node status) stages, pathological tumor size, histological type and number of metastatic lymph nodes. In addition tissue microarray array analysis was performed from the tumor samples. Area percentage of staining was determined using immunohistochemistry for estrogen receptor (*ER*), progesterone receptor *(PR), HER2, Ki67, CK56, EGFR, VIM, CD24, CD44, ALDH1, BCL2*, E-Cadherin and Trio^25^. Tumors were classified as *HER2* positive if the *HER2* immunostaining showed 3+ intensity or from individual review for 2+ scores^25^.

Missing covariate values were imputed before model building using the *missForest* imputation algorithm^27^. The percentage of missing data was less than 5% for all variables (Figure S1). Using 100 trees per forest (predefined setting value of *missForest)*, continuous and categorical covariates were imputed with a 4.4% and 7.1% error, respectively.

Institutional review board approval was obtained for this retrospective study in accordance with national laws.

### Random survival forests analysis (RSF)

The RSF algorithm is an extension of Breiman’s random forest for the analysis of right-censored time-to-event data^28^. We utilized the RSF implementation of the randomForestSRC *R* package^29^. All RSF models were fitted using 1000 trees, with the logrank splitting rule^28^. The optimal values of the tuning parameters (number of variables to be sampled at each split and minimum number of data points in a terminal node) were selected to maximize the concordance index calculated on the out-of-bag data.

Impact of covariates on the TTR was assessed using the forest-averaged minimal depth^30^, which quantifies the predictive value of a covariate in a tree by its distance from the root node to the first node where it is used to split (smaller minimal depth values correspond to more predictive covariates).

### Mechanistic model of metastatic dissemination and growth

The individual primary tumor (PT) kinetics in individual *i* were described by the Gompertz model:

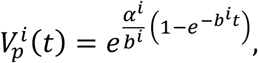

where 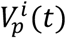 is the number of cells within the PT at time *t* and *α^i^* and *b^i^* are the Gompertzian growth parameters. Under this model, the tumor will approach a theoretical upper limit 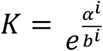. In order to avoid overfitting and improve identifiability, *K* was fixed to 10^12^ cells, leaving *α^i^* as the only free parameter driving growth. This value was used following statistical estimations and biological considerations in breast cancer^18, 31^. The PT size - reported as a diameter in the data - was converted into number of cells assuming spherical shape and the assumption 1mm@ = 10^6^ cells^31, 32^. All model simulations were performed in number of cells.

Considering a dissemination rate from the PT given by^22^

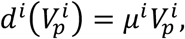

the total number of metastases at time *t* is

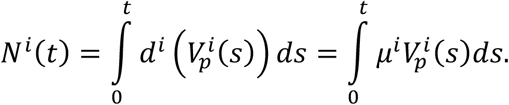

The individual parameter *μ^i^* is the per day probability for a PT cell to disseminate and establish a distant metastatic colony. Each metastasis was assumed to start from the volume *V*_0_ of a single cell and to grow at the same rate than the PT:

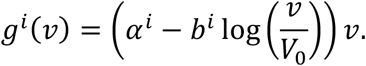

The state of the metastatic process was described by a function *ρ^i^*(*t,v*) representing the distribution of metastatic tumors with size *v* at time *t*. It is given by the solution of the following transport equation^21^:

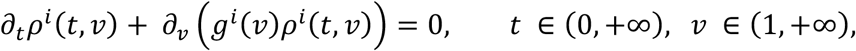

endowed with the boundary and initial conditions:

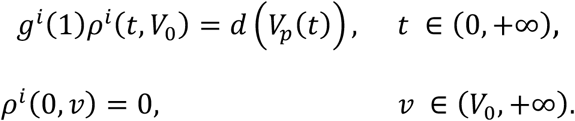

### Mechanistic modeling of the time-to-relapse

To calibrate the metastatic model on TTR data, we defined the theoretical time to relapse as illustrated in Figure 1. More precisely, assuming a value *V_vis_* as detection threshold, the time *τ_vis_* for a tumor to reach this size was given from the assumption of Gompertzian growth, i.e.

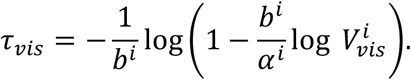

**Figure 1:**
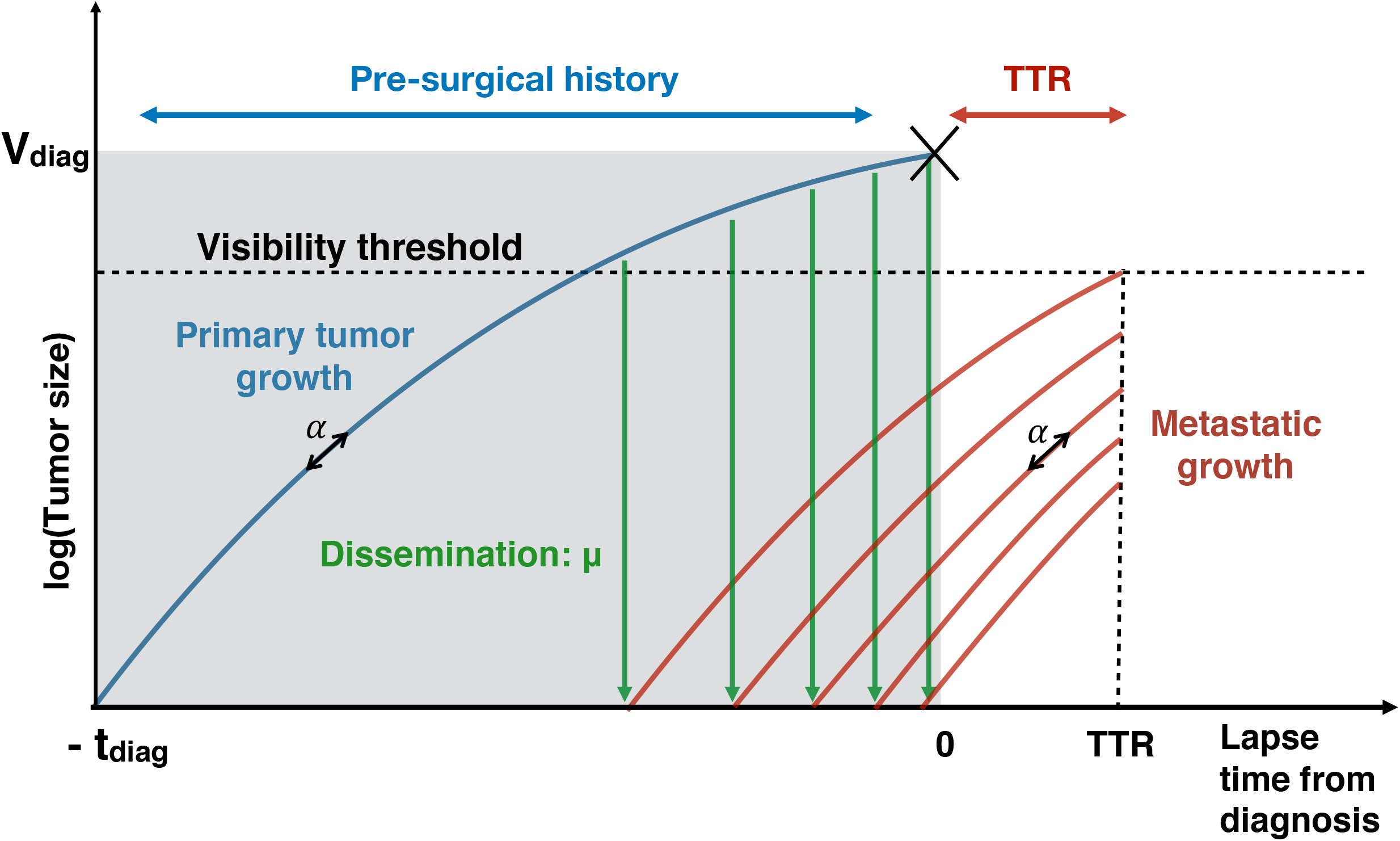
Scheme of the mechanistic model. Growth of primary and metastatic tumours are characterized by a common growth parameter *α*. Emission of metastases from the primary tumour occurs at a rate that depends on the primary tumour size and on a parameter of metastatic potential *μ*. The time-to- recurrence is defined as the lapse time from diagnosis until the first emitted metastasis reaches the visible size.

In an analogous way, the time from the first cancer cell to the detection of the primary tumor, 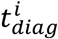, was determined from the known size of the primary tumor at detection, 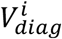:

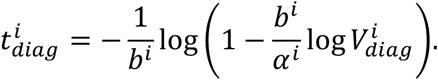

A visibility threshold *V_vis_* of 5 mm in diameter was assumed to be the detectability limit at imaging.

Since metastases of size larger than *V_vis_* at time *t* must have been emitted in the time interval 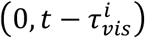, the number of visible metastases at time *t* can be obtained by

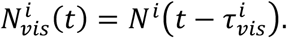

The theoretical TTR was then defined as

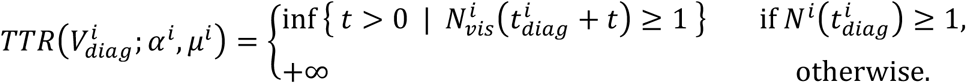

That is, the time elapsed from diagnosis to the appearance of the first visible metastasis, if, according to the model, at least a metastasis was emitted before diagnosis; otherwise it was considered as infinite.

### Calibration of the mechanistic model using mixed-effects learning

The mathematical model for individual TTR was thus defined as a function of 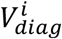 and a vector of structural individual parameters *ψ^i^* = (*α^i^,μ^i^*). Let *T^i^* denote the TTR for patient *i*. A constant error model was assumed on the log-transformed data:

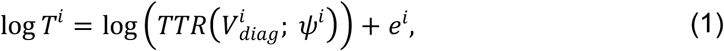

where *e^i^* is the residual error following a normal distribution with mean 0 and variance *σ*^2^. The log-transformation was used to ensure positive values of the TTR variable. The individual parameters were assumed to be log-normally distributed and a linear covariate model was used:

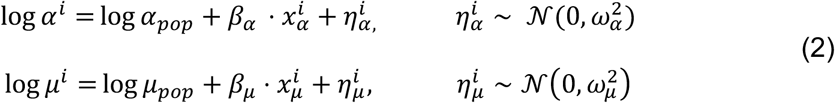

where 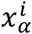, and 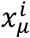 are vectors of subject-specific covariates, which might be identical, or partially or completely different.

The statistical model for the observations (1) implicitly defines for each individual *i* the probability density function of *T^i^* conditionally on the individual parameters *ψ^i^*. The corresponding survival and hazard functions can be obtained by:

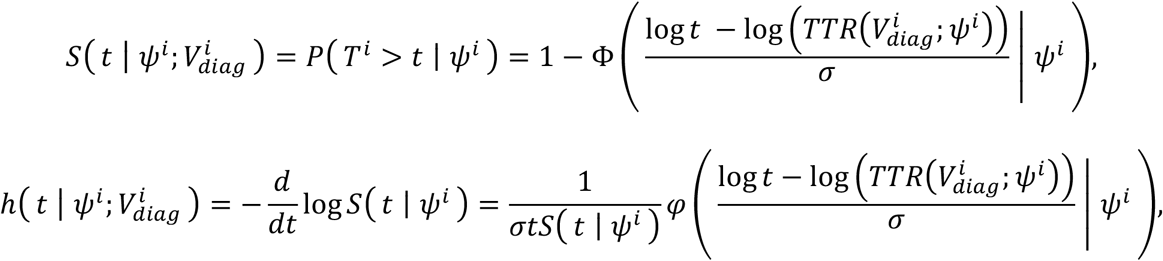

where Φ and *φ* are the cumulative distribution and probability density functions of the standard normal distribution, respectively.

Let *C^i^* denote the time to death or last follow-up for individual *i*. With right-censoring the observations are 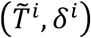, where 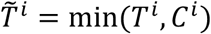 and 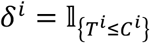 is the indicator variable. The contribution of individual *i* to the likelihood is then given by^33^:

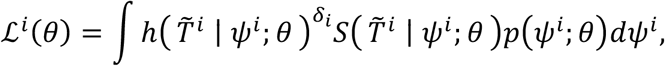

where *θ* = (*α_pop_, μ_pop_, β_α_ β_μ_,ω_α_,ω_μ_,σ*) are the population parameters to be estimated and *p*(*ψ^i^; θ*) is the probability density function of the individual parameters, defined by (2).

The likelihood function (product of the individual likelihoods 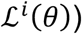) was maximized using the Stochastic Approximation of Expectation-Maximization (SAEM) algorithm^24^ implemented in the *R* saemix package^34^. Initial values of the parameters were *α*^0^ = 0.002, *μ*^0^ = 4 × 10^-12^, 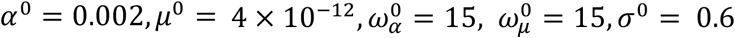. Standard errors of the estimated parameters were obtained using 100 bootstrap samples, and significance of covariates was assessed using the Wald test.

The mean of the conditional survival functions was estimated as 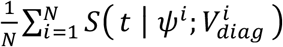, with (*ψ*^1^,…,*ψ^N^*) and 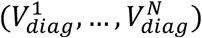 drawn respectively from the estimated distribution of the individual parameters and the values of *V_diag_* in the data.

### Classification machine learning algorithms for prediction of 5-year distant metastasis-free survival

Machine learning algorithms were trained using the scikit-learn python package^35^. These included logistic regression, support vector machines (SVM), random forests (RF), k-nearest neighbors (kNN) and gradient boosting. Values of the models hyperparameters were selected to optimize the area under the ROC curve, using a 3-fold cross-validated grid search. The kernel used for SVM was the radial basis function *K*(*x,x′*) = *e*^-*γ*||*x-x*′||^ with 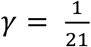 (21=number of features) and regularization parameter *C* = 1. For the classification algorithms (and for them only), because these are not able to manage censored data, patients with data censored before 5 years were removed from the dataset, leaving a total of 594 patients. A balanced version of the dataset was constructed by keeping all 55 patients with 5-years metastatic relapse and by adding an equal number of patients randomly selected from those who experienced metastatic relapse later than 5 years from diagnosis. Each model was evaluated using 100 replicates of 10-fold cross-validations, with folds created using a stratified random sampling strategy (to preserve class balance). We applied a standard scaling for each run and each set by removing the training folds mean and scaling to unit variance, for the algorithms that require a homogeneous scale (SVM, kNN). The random forests algorithm was also seeded with a different random initialization value each time.

### Evaluation of predictive performances

All models were internally validated using 10-fold cross-validation. For classification tasks, the models were evaluated using receiver operating characteristic (ROC) curve analysis and standard performance metrics for binary classification algorithms^36^.

Discrimination of time-to-event models was quantified by Harrell’s c-index^37^. Calibration of each model was examined graphically, by comparing the model predicted probabilities against the observed event rates^37^.

## Results

### Random survival forest multivariate analysis

For machine learning analysis of the TTR data, we used the RSF algorithm, which allows right-censored data^30^. In addition to evaluating its predictive power, we used this algorithm to identify variables most predictive of TTR. Covariates were ordered on the basis of minimal depth (Figure 2A) and selected by running a nested analysis (Figure 2B). The crossvalidated c-index improved as the number of variables increased, reaching a maximum of 0.67 (95% CI, 0.66-0.69) with 5 variables. We selected these 5 top variables for the optimal model and future analysis in the mechanistic model below. Calibration plots for 2-, 5- and 10-year demonstrated a good agreement between the observed and model-predicted probabilities of metastatic relapse (Figure 2C), with moderate overprediction of relapse in the higher risk groups.

**Figure 2:**
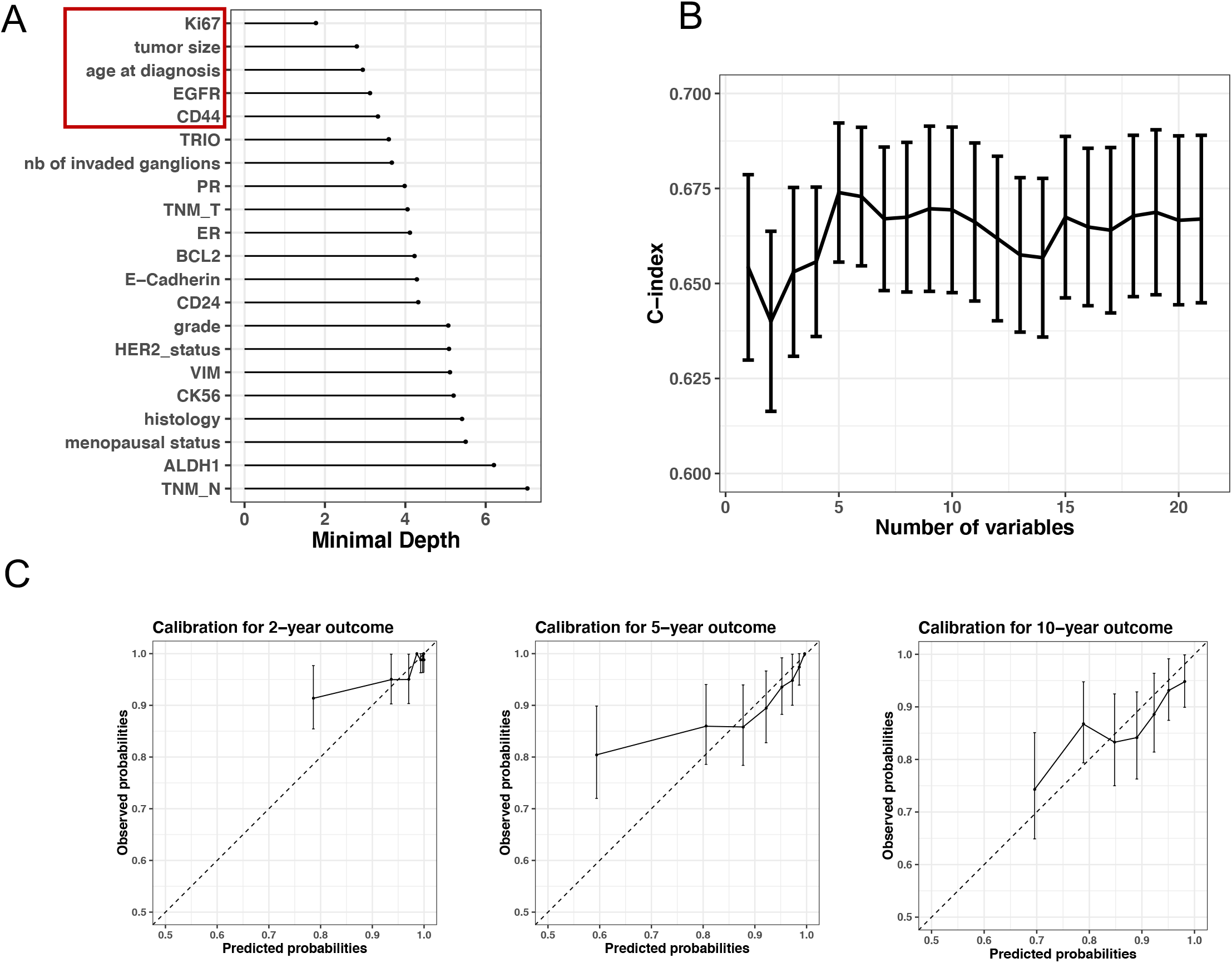
Survival random forest and covariate selection. A) Minimal depth ranking of covariates. B) Cross-validated Harrel’s c-index under sequentially fit RSF models with variables ordered by importance using minimal depth. Bars are 95% confidence intervals. C) Calibration plots for the RSF model with the 5 top variables (red box in figure A). Bars are 95% confidence intervals.

Partial dependence plots for the selected covariates indicated strong nonlinear relationships between covariates and relapse probabilities, with a non-monotonic behavior for age and tumor size (Figure S2). Confirming previous results^38^, these plots suggest nonvalidity of the proportional hazards assumption. Indeed, if such assumption would hold, the probability of no relapse at any time would be monotonous as a function of any covariate value.

### Calibration and validation of the mechanistic model

To offer better insights on the mechanisms of relapse, we developed a mechanistic model of the TTR. We first evaluated the ability of this model to describe the TTR data without using covariates, except for pathological tumor size, which is a variable encoded in the structural model. We asked whether we could describe inter-individual variability of TTR by means of population statistical distributions of the parameters *μ* (dissemination) and *α* (growth) of the model. Estimates of the population parameters were obtained using the SAEM algorithm and are reported in Table 1. Both fixed and random effects were identified with satisfactory precision (relative standard error < 37%). Figure 3A compares the model estimation of the TTR survival function to the empirical Kaplan-Meier estimate. Despite a slight overestimation of the metastatic risk for shorter times, the model was able to capture the shape of the Kaplan-Meier estimate. To further verify the agreement between model and data, we also compared model and Kaplan-Meier curves for different values of the tumor size at diagnosis (Figure S3). Although the model remained within the Kaplan-Meier confidence interval in all cases, we observed that model predictions tended to be less accurate as the tumor size increased, with overestimation of the relapse risk for large PT sizes.

**Figure 3:**
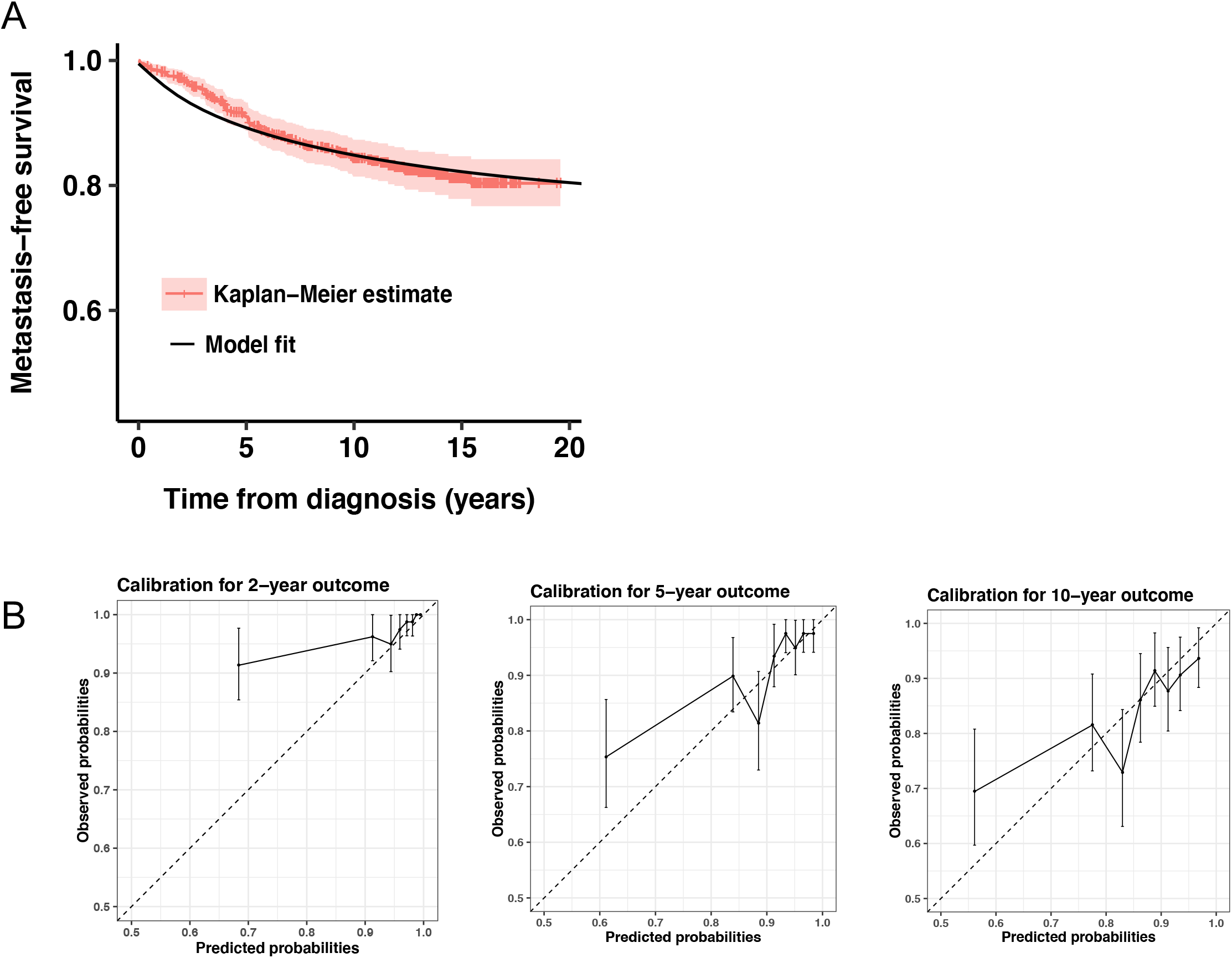
Fit of the mechanistic model. A) Mean of the conditional metastasis-free survival functions from the mechanistic model and Kaplan-Meier estimates with 95% confidence interval. B) Calibration plots of metastasis-free survival for the final mechanistic model with covariates.

**Table 1:** Parameter estimates. Mechanistic mixed-effects models for the time to metastatic relapse were fitted to the data using likelihood maximization. Parameters *α* (growth) and *μ* (dissemination) were assumed to follow lognormal distributions with fixed effects (typical value) *a_pop_* and *μ_pop_*, standard deviation of the random effects *ω_α_* and *ω_μ_* and standard deviation of the error model *σ* (see methods). A first version was fitted without covariates (top) and a second with inclusion of dependence of the individual parameters on individual covariates (bottom). r.s.e.: relative standard error. p-value refers to a Wald test for statistical significance of the covariate on the parameter value.

|  | Parameter | Estimate | r.s.e. (%) | p-value |
| --- | --- | --- | --- | --- |
| Model without<br>covariates | $\log \alpha_{pop}$ | -6.34 | 12.6 | |
| | $\log \mu_{pop}$ | -26.8 | 3.68 | |
| | $\sigma$ | 0.542 | 28.4 | |
| | $\omega_{\alpha}$ | 3.37 | 36.4 | |
| | $\omega_{\mu}$ | 3.78 | 15.9 | |
| Model with<br>covariates | $\log \alpha_{pop}$ | -9.01 | 10.8 | |
| | $\beta_{\text{Ki67},\alpha}$ | 0.093 | 29.6 | 0.001 |
| | $\beta_{\text{CD44},\alpha}$ | 0.017 | 57.7 | 0.083 |
| | $\log \mu_{pop}$ | -25.9 | 4.4 | |
| | $\beta_{\text{EGFR},\mu}$ | 0.053 | 38.1 | 0.009 |
| | $\sigma$ | 0.606 | 24 | |
| | $\omega_{\alpha}$ | 2.75 | 22.1 | |
| | $\omega_{\mu}$ | 3.03 | 20.5 | |

### Mechanistic covariate analysis and predictive power of the mathematical model

We next tested the covariates selected by the RSF analysis in the mechanistic model. We built the covariate model using a backward elimination procedure, starting with the full model with all the preselected covariates on both parameters *α* and *μ*. The final model included *Ki67* and *CD44* on *α*, and *EGFR* on the dissemination parameter *μ* (Table 1). The crossvalidated c-index for this model was 0.65 (95% CI, 0.60-0.71). Calibration plots for 2-, 5- and 10-year outcomes demonstrated good predictive accuracy of the model (Figure 3B). Nevertheless, similarly as the RSF model, risk of relapse was overestimated in high risk groups. For the classification task of prediction of 5- and 10-year relapse, comparable results were obtained between the mechanistic model, the RSF, classification machine learning algorithms and Cox regression (Table 2).

**Table 2:** Prediction metrics for classification of 5- and 10-years metastatic relapse. For comparison purpose between time-to-event and classification models, prediction metrics performed on the entire data set are reported. In bold is the best score achieved for a given metric.

|  | Algorithm | AUROC | Accuracy | Sensitivity | Specificity | PPV | NPV | F1 |
| --- | --- | --- | --- | --- | --- | --- | --- | --- |
| 5 years | *Mechanistic model* | *0.73* | *0.68* | **0.75** | *0.67* | *0.19* | **0.96** | *0.30* |
|  | Random survival forest | 0.73 | 0.69 | 0.64 | 0.70 | 0.18 | 0.95 | 0.28 |
|  | Random forest | **0.75** | 0.66 | 0.71 | 0.66 | 0.18 | **0.96** | 0.28 |
|  | Logistic regression | **0.75** | 0.83 | 0.42 | 0.87 | 0.24 | 0.94 | **0.31** |
|  | k-nearest neighbor | 0.62 | **0.91** | 0.02 | **1.00** | **0.41** | 0.91 | 0.05 |
|  | Gradient boosting | 0.71 | 0.90 | 0.11 | 0.98 | 0.36 | 0.92 | 0.17 |
|  | Support vector machine | 0.64 | 0.87 | 0.09 | 0.95 | 0.15 | 0.91 | 0.11 |
|  | Cox | 0.71 | 0.72 | 0.66 | 0.73 | 0.20 | 0.95 | **0.31** |
| 10 years | *Mechanistic model* | *0.67* | **0.67** | *0.62* | **0.68** | **0.30** | *0.89* | **0.41** |
|  | Random survival forest | **0.69** | 0.62 | **0.71** | 0.60 | 0.28 | **0.90** | **0.41** |
|  | Cox | 0.65 | 0.65 | 0.61 | 0.65 | 0.28 | 0.88 | 0.39 |

### Predictive simulations of the mechanistic model

The previous results allow to calibrate the mechanistic model parameters from variables available at diagnosis, by using only the covariate part in equation (2) and neglecting the remaining unexplained variance. We used this to simulate the natural cancer history for a number of representative patients of our dataset. For each patient, the population-level parameters *α_pop_,μ_pop_,β*_*κi*67,*α*_,*β*_*CD*44,*α*_ and *β_EGFR,μ_* were calibrated from an independent training set that did not contain this patient (coming from the cross-validation procedure). Simulations were then performed using a discrete version of the metastatic model^22^ and are reported in Figures 4 and S4. Covariate values used for prediction and resulting inferred individual parameters 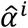 and 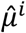 are reported in Table S1.

**Figure 4:**
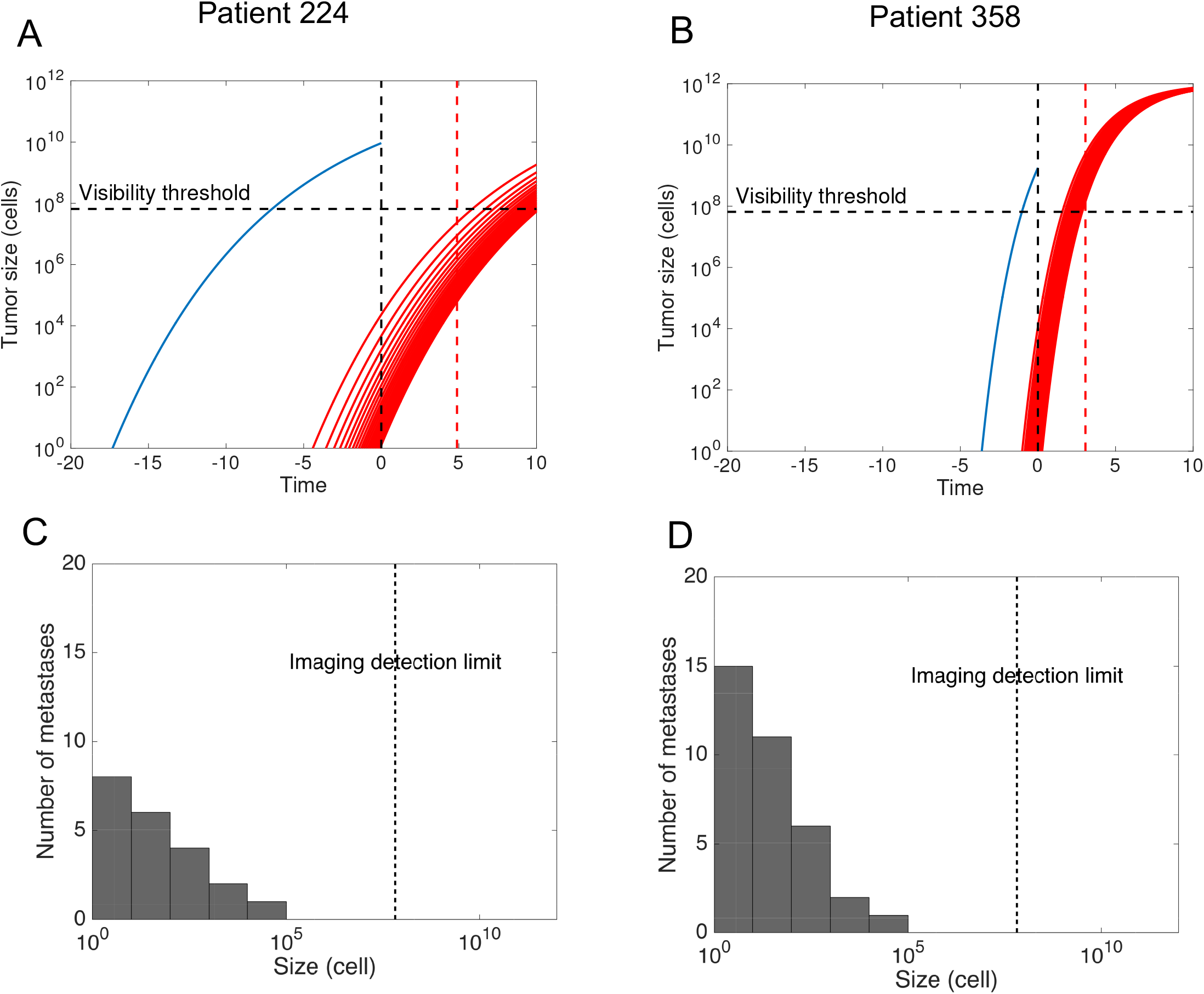
Simulations of the cancer history for two patients. Simulations were performed using predicted values of the individual parameters 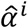 and 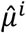 as 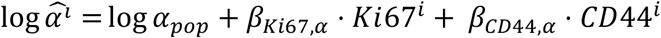. and 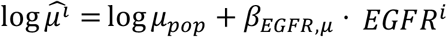. Population parameters *α_pop_, β_Ki67,α_,β*_*CD*44,*α*_, *μ_pop_* and *β_EGFR,μ_* were inferred from a training set that did not contain patient *i*, during the cross-validation procedure. A-B): Growth of the primary tumor and metastases that will ultimately become visible at a 10 years post-surgical horizon. The red dashed line indicates the observed time of metastatic relapse C-D): Size distribution of the invisible metastases at the time of diagnosis.

For patients 224 and 358 (Figure 4A-B), the model predicted a time from the first cell to detection (pre-surgical history) of 17.3 years and 3.61 years, respectively. In addition, the model allowed to predict the metastatic size distribution at diagnosis (Figure 4C-D). Patient 224 was predicted to have 21 invisible metastases at the time of diagnosis. The largest metastasis contained 2.32 · 10^4^ cells and the smallest only 1 cell. The former was predicted to have initiated 12.9 years after the first cell of the PT, i.e. 4.4 years before surgery. The model predicts relapse 6.18 years after diagnosis while true relapse occurred at 4.88 years. At diagnosis, patient 358 was predicted to have a total of 35 invisible metastases. The largest metastasis contained 1.16 · 10^4^ cells and the smallest 1.36 cells. According to this simulation, the first metastasis in this patient was emitted 2.6 years after the PT onset, i.e. 1.01 years before surgery. The model predicted relapse 1.97 years after diagnosis while true relapse occurred at 3.06 years. The differences in PT and metastatic dynamics for these two patients are due to the different values of the covariates (Table S1). Distinct levels of *Ki67* cause distinct growth kinetics. Moreover, unlike patient 224, the tumor of patient 358 expresses *EGFR*, which is associated with a higher metastatic potential. Thus although patient 358’s tumor is much younger, the total number of (invisible) metastases at surgery is predicted to be larger (35 vs 21 metastases in patient 224). Model predictions are also informative in the case of individuals who were censored at the last follow-up. For instance, patient 70 (Figure S4) was censored at 17.7 years after diagnosis. This is consistent with our model, which predicts that this patient was disease-free after PT resection and would never have relapsed (TTR = +∞).

### Comparison with machine learning classification algorithms and Cox regression

We tested the predictive power of machine learning classification algorithms. These cannot account for right-censored data. Thus, for this part we focused on prediction of 5-years metastatic relapse (yes or no). Best performances were achieved by the random forest and logistic regression models (Table 2 and Figure S5A-D). However, owing to the low event rate (9.25%), positive predictive value and and F1 scores were low (Table 2). To improve these metrics, a balanced, downsampled version of the dataset was constructed. This significantly improved PPV and F1, as well as model calibration (Figure S5E-F). We also compared our results to classical Cox regression survival analysis, which was found to exhibit similar predictive power (Figure S6, Tables 2 and S2).

## Discussion

We propose a mechanistic model for prediction of metastatic relapse after surgical intervention in patients with early-stage breast cancer which, for the first time, is able to simulate the pre- and post-diagnosis history of the disease from data available at diagnosis only. Notably, with a c-index of 0.65 (95% CI, 0.60 - 0.71), the mechanistic model achieved similar performances as the RSF algorithm (95% CI, 0.66-0.69) and Cox regression (95% CI, 0.62-0.67). These were also comparable with the actual standard prognosis model of breast-cancer-specific survival Adjuvant! (c-index 0.71)^7^, used for risk classification in the MINDACT trial^39^. Others also reported a similar c-index of 0.67 for prediction of relapse in untreated patients, using Cox analysis^40^. Advanced deep learning algorithms did not outperform this predictive power (c-index 0.68) for prediction of survival, even though integrating genomic data^13^. A recent study that considered recurrence (either local, regional or distant) reported superior predictive power (AUC 0.81 for prediction of relapse at 5-years, versus 0.73 in our analysis), which might be explained by the much larger data set (15, 314 patients) and inclusion of epidemiological data not available in our analysis.

Grounded on the biology of the metastatic process, our model provides insights not achievable by statistical analysis alone. First, one of the most important and well known predictor of metastatic relapse - tumor size^23^ - is directly incorporated into the model as an input parameter, used for calculation of the tumor age. Second, our model allows to test whether covariates are associated with growth and/or dissemination. We found that *Ki67* was positively associated with the proliferation parameter *α*, which aligns with the definition of *Ki67* as a proliferation marker^41^, as well as other results confirming its predictive power^42^. On the other hand, the basal marker *EGFR* was found to be a significantly positive covariate for the dissemination parameter *μ. EGFR* has previously been demonstrated as being a prognostic factor of relapse^43^, yet it is interesting to note that correlation appears here on *μ*, which is consistent with the known fact that basal-like breast cancers are metastatically more aggressive^44^. Interestingly, the median value of *μ* was consistent with the value estimated in a previous work using data of metastatic relapse probabilities from a cohort of breast cancer patients (*μ_pop_* = 2.26 × 10^-12^ cell^-1^.day^-1^ here vs *μ_pop_* = 7 × 10^-12^ cell^-1^.day^-1^ in^22^).

Our analysis also confirmed the prognostic value of age at diagnosis, with younger patients having a higher risk of relapse^45^. However, we found that after 60 years-old, the risk of relapse was increasing (Figure S2). This nonlinearity might explain why age did not appear as significant in either our mechanistic or Cox analysis. Although not significant at the 0.05 threshold, our results suggest prognosis value of *CD44* (p = 0.083 for association with *α*). *CD44* is a cellular protein which is used as a marker of breast cancer stem cells^46^ and is associated with metastasis^47, 48^. Since *CD44* is involved in cell adhesion and invasion^49^, it is surprising that it emerges as associated with *α* and not *μ*. This might indicate limitations of our model to detect complex biological processes since inference is only made indirectly and using several simplifying assumptions, such as a unimodal lognormal distribution of the parameters in the population. This might also be due to the fact that we considered here *CD44* as a continuous variable, whereas a standard marker is the percentage of *CD44^+^/CD24^-^* cells^46^. Similarly, only *EGFR* as a continuous variable was investigated and not the more classical *EGFR^+^/CK5/6^+^* marker^43^. Indeed, the aim of the current study was to establish the methodology of using our mechanistic model as a predictive tool, and we favored first keeping continuous variables. We plan to perform more detailed examination of such clinical variables in forthcoming work, as well as study the predictive power of the model in well-established subgroups such as node-negative patients or patients stratified according to the current molecular classification^50^.

A major advantage and clinical relevance of the mechanistic model over standard statistical or machine learning models, is that it can be used to perform patient-specific simulations allowing to assess the extent of invisible metastases at the time of diagnosis and to predict future growth of metastases. In turn, this might aid selecting patients that will most benefit from extended adjuvant therapy (or conversely, patients who would need only a limited number of cycles), by performing individualized simulations of the future course of the disease under competing therapeutic strategies. However, it will be first required to develop and validate models integrating the effect of systemic adjuvant therapies^51^. We also believe that our methodology could be applied to other cancer types where similar concerns occur about the use of adjuvant therapy to avoid metastatic relapse (e.g. lung or kidney cancer). In addition, the novel approach we propose to mechanistically model time-to-event data could be used to extract biologically relevant information from such data, which - although ubiquitous in clinical oncology - are almost exclusively analyzed using agnostic statistical tools.

Our model represents a first attempt of a mechanistic, individual-level, predictive model of metastatic relapse and might be improved in a number of ways. For instance, unexplained variability remained important despite the inclusion of covariates, suggesting that biomarkers other than those tested might improve model predictions. In this regard, genetic expression signatures have been shown to have higher predictive power compared to standard histological and clinical variables alone^52^. Our mechanistic model could also be refined by higher order phenomena such as dormancy, which has been proposed to explain recurrence occurring after many years from surgery^53, 54^. Finally, to be applied in clinical practice, the model should be further evaluated on external data sets.

## Acknowledgments

The authors thank Institut Bergonié for permission to use the data from the Breast Cancer Database for this research. They also thank the “Big bang” association for funding support, in memory of Marie-Christine Masini.

## Supplementary material

**Figure S1:**
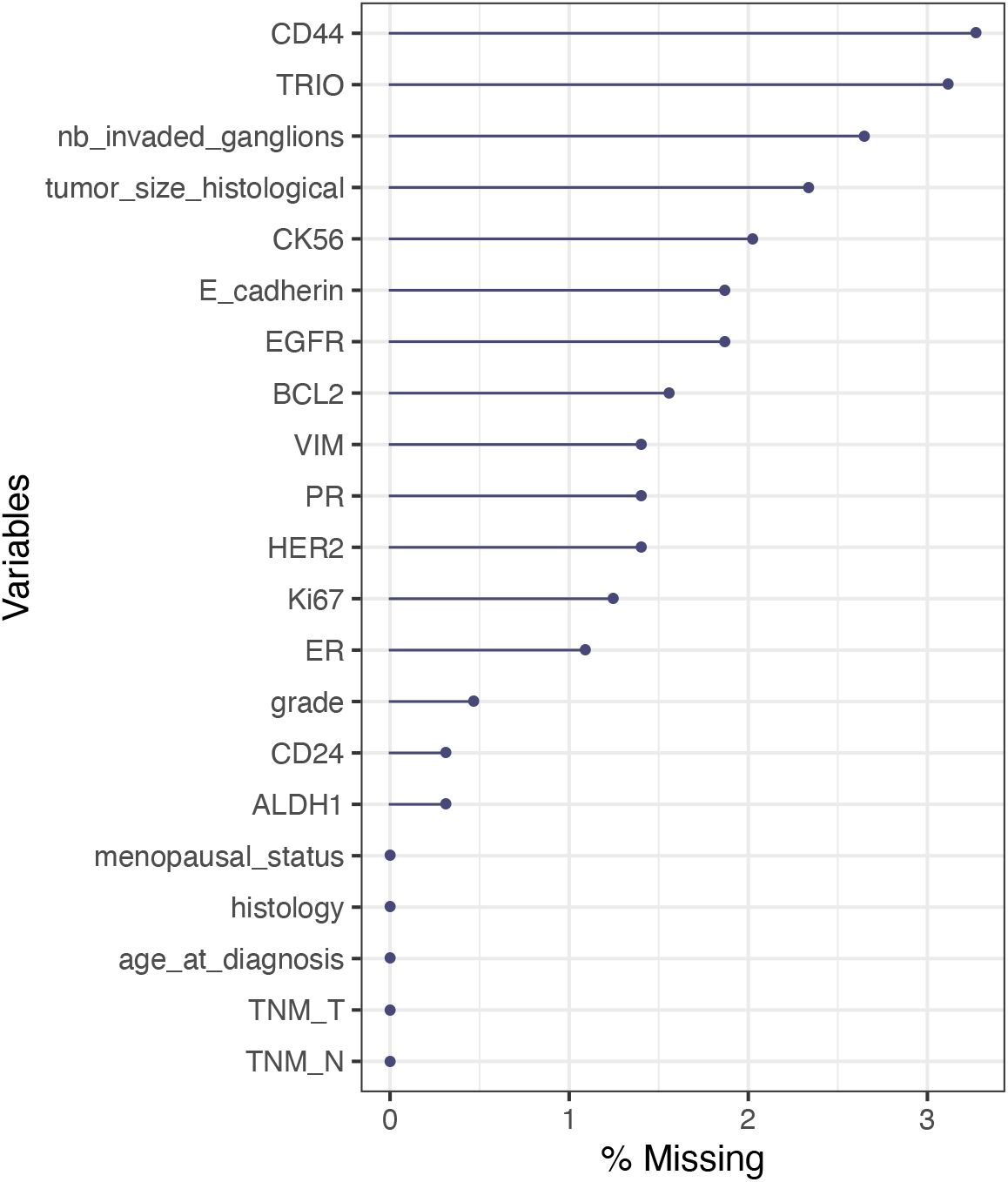
Percentage of missing values in each variable

**Figure S2:**
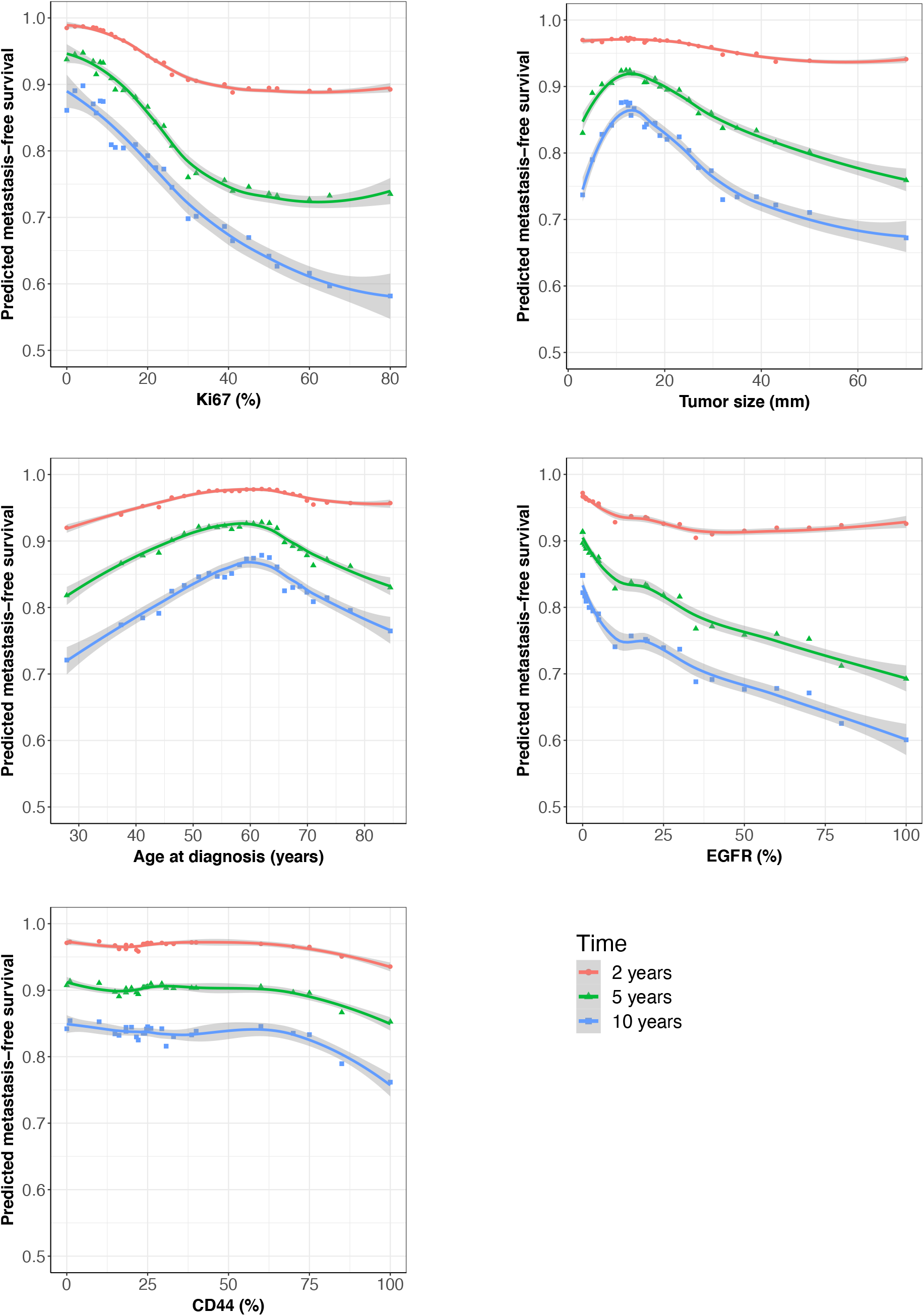
Partial dependence plots of the random survival forest predicted DMFS as a function of the top eight predictors according to the minimal depth ranking

**Figure S3:**
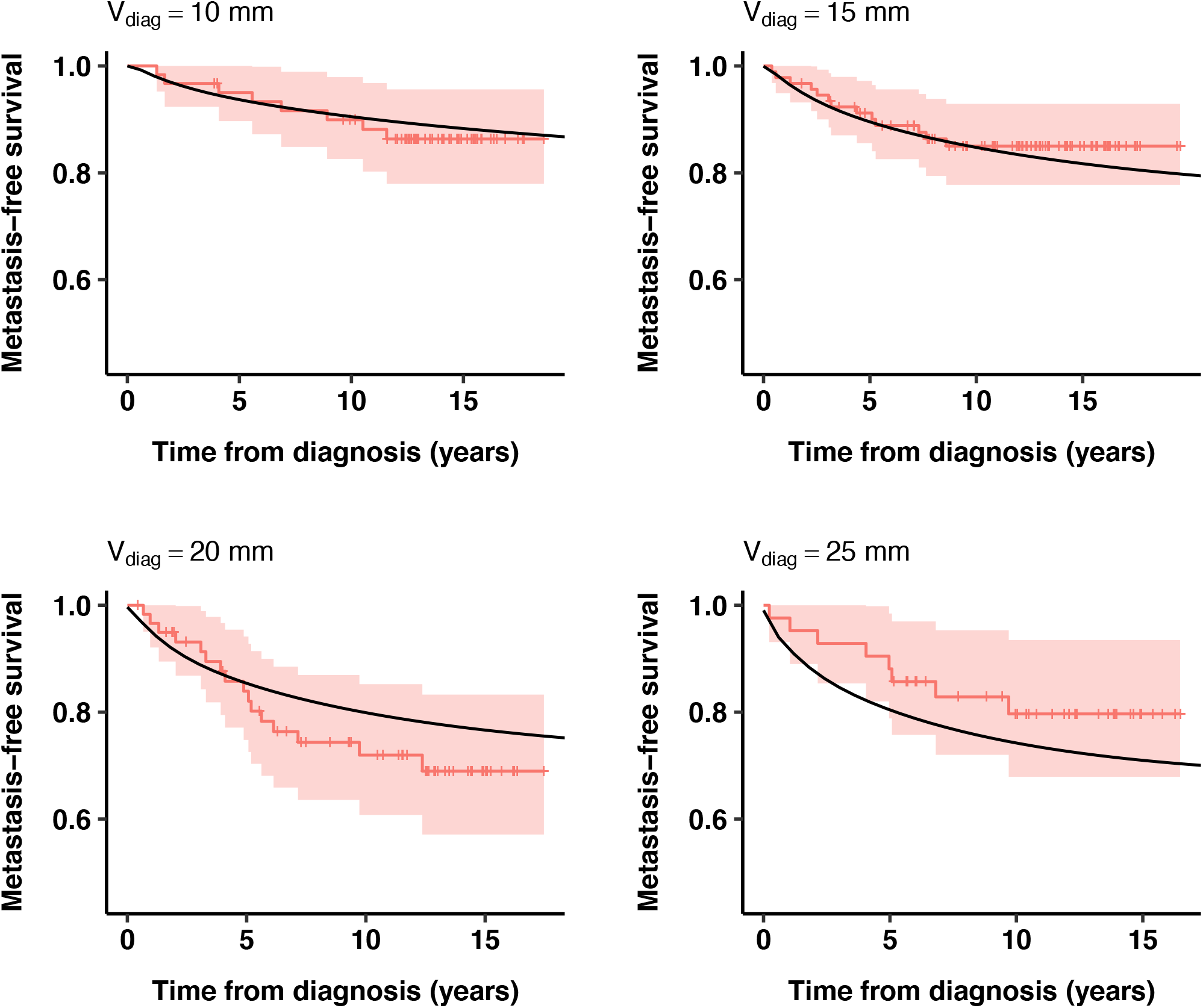
Distant metastasis-free survival predictions of the mechanistic model and Kaplan-Meier estimates with 95% confidence intervals stratified by values of the primary tumour size (diameter) at diagnosis

**Figure S4:**
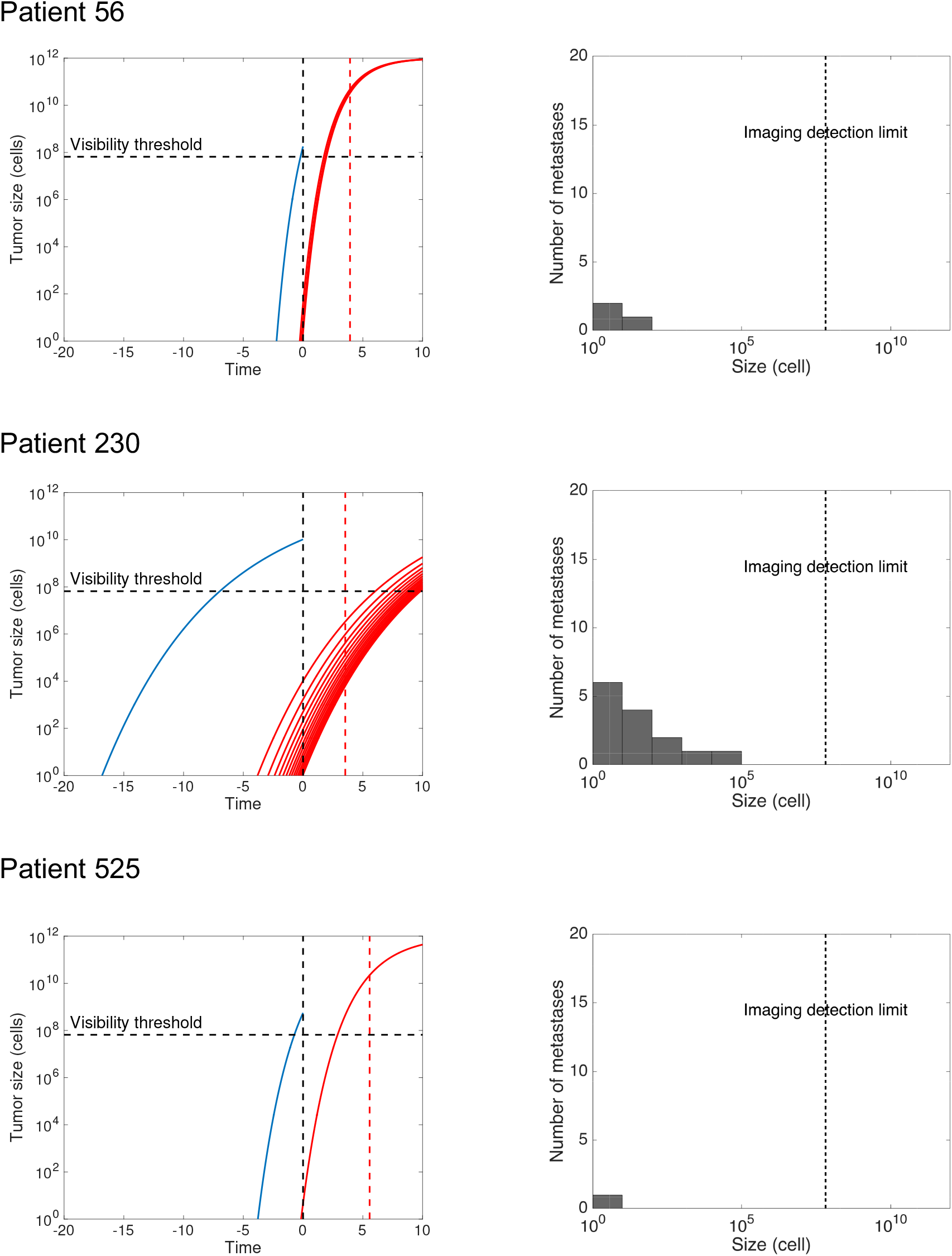
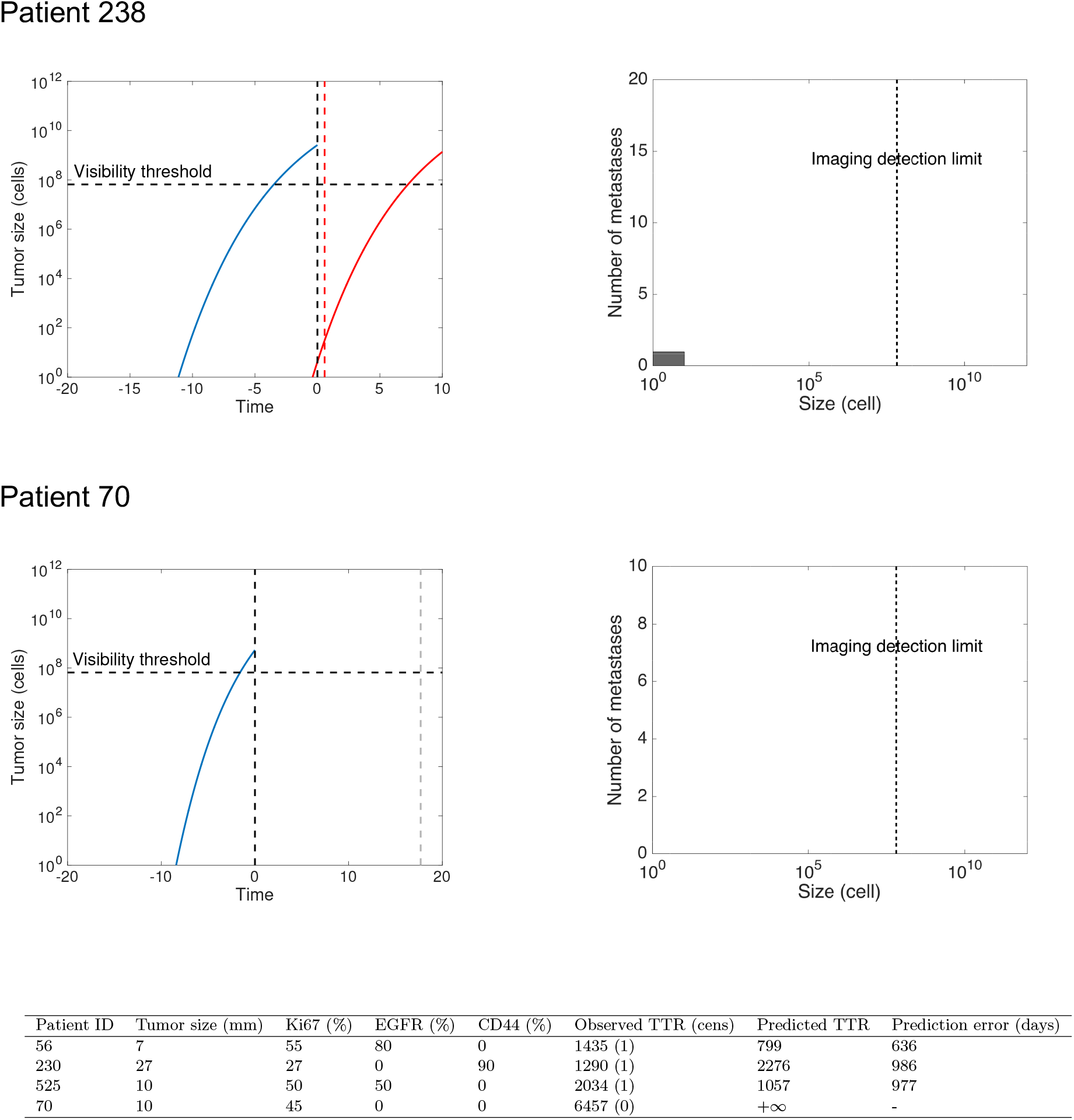
Predictions of the mechanistic model for individual patients

**Figure S5:**
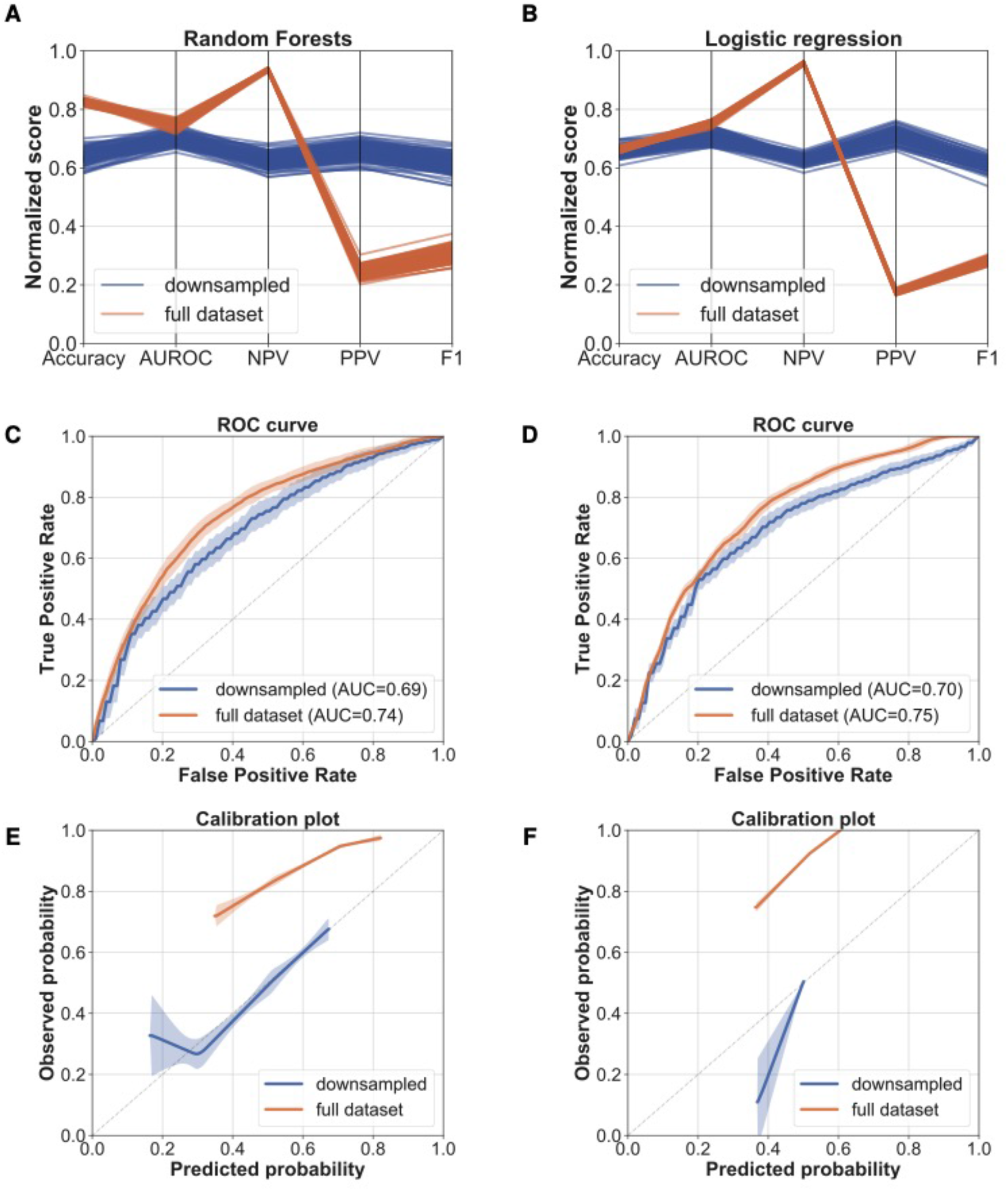
Prediction of 5-years metastatic relapse using machine learning classification algorithms. *Left:* Metrics and plots for the random forests and logistic regression algorithms for the full dataset (brown) and the downsampled dataset (blue). A-B) Metrics parallel plot representing scores of accuracy, area under the receiver-operator curve (AUROC), negative predictive value (NPV), positive predictive value (PPV) and F1 (100 replicates). C-D) ROC curves. Mean ± entire range (100 replicates). E-F) Calibration plots. Mean ± entire range (100 replicates).

**Figure S6:**
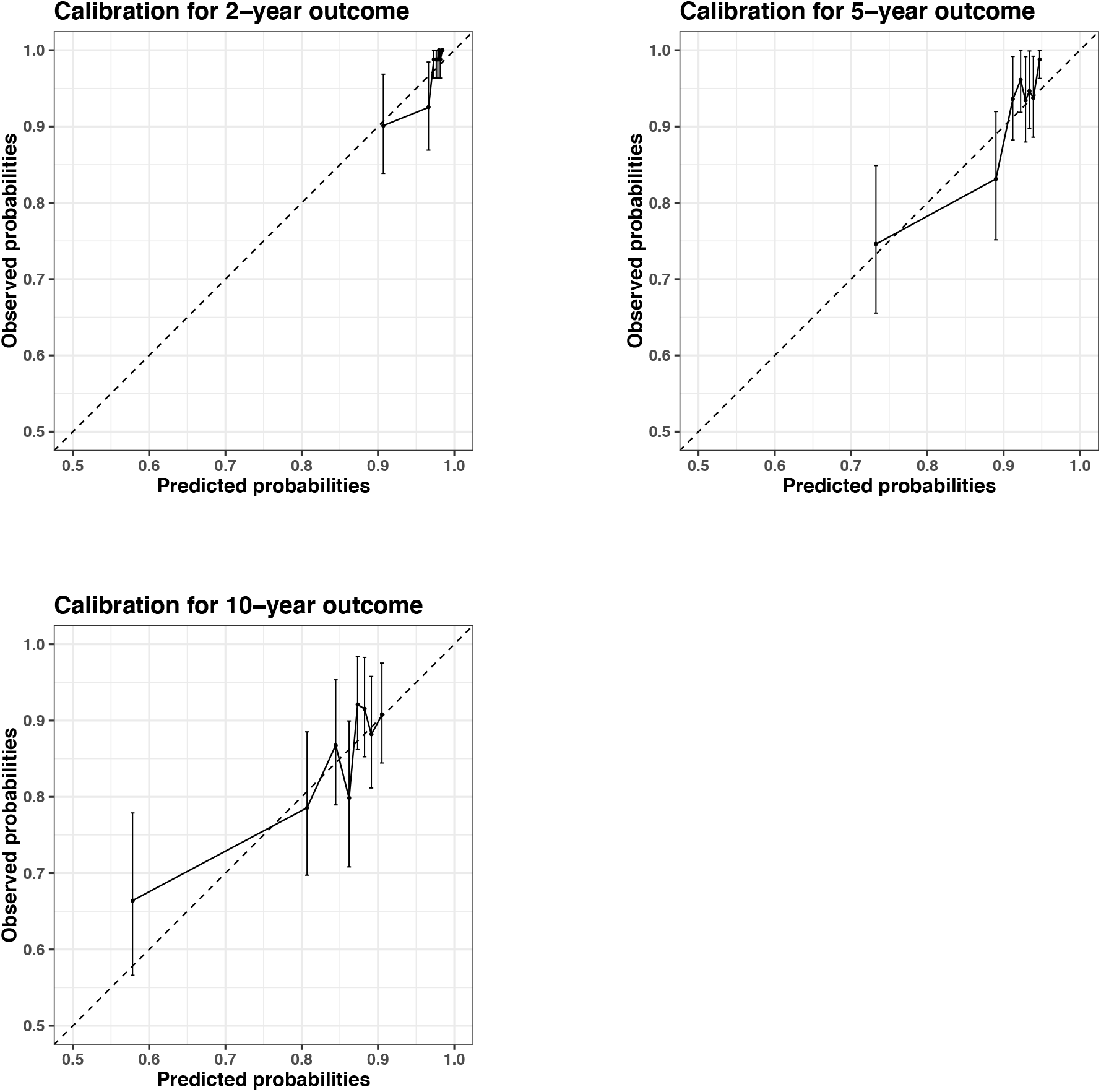
Calibration plots for the Cox model with the eight covariates selected through the RSF analysis

**Table S1:**
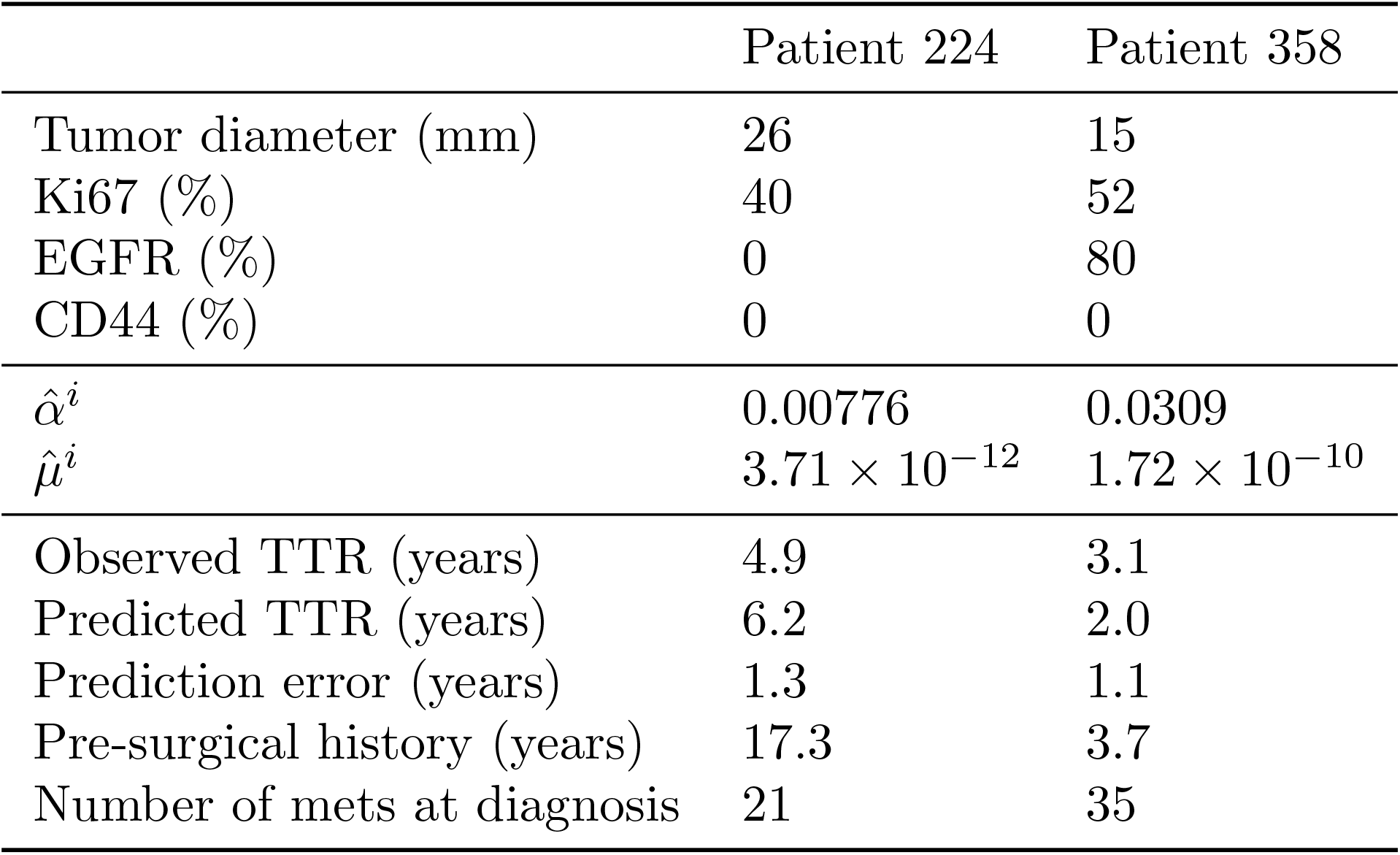
Covariates and resulting inferred parameter values for individual predictions. TTR = time-to-relapse

|  | Patient 224 | Patient 358 |
| --- | --- | --- |
| Tumor diameter (mm) | 26 | 15 |
| Ki67 (%) | 40 | 52 |
| EGFR (%) | 0 | 80 |
| CD44 (%) | 0 | 0 |
| $\hat{\alpha}^i$ | 0.00776 | 0.0309 |
| $\hat{\mu}^i$ | $3.71 \times 10^{-12}$ | $1.72 \times 10^{-10}$ |
| Observed TTR (years) | 4.9 | 3.1 |
| Predicted TTR (years) | 6.2 | 2.0 |
| Prediction error (years) | 1.3 | 1.1 |
| Pre-surgical history (years) | 17.3 | 3.7 |
| Number of mets at diagnosis | 21 | 35 |

**Table S2:** Cox regression using the first five covariates selected by minimal depth with the random survival forest model. HR = hazard ratio. 95% CI = 95% confidence interval.

|  | HR | 95% CI | p-value |
| --- | --- | --- | --- |
| Ki67 | 1.02 | [1.01, 1.03] | $1.8 \cdot 10^{-5}$ |
| Tumor size | 1.01 | [0.99, 1.03] | 0.33 |
| Age | 0.99 | [0.98, 1.01] | 0.43 |
| EGFR | 1.01 | [1.00, 1.02] | 0.04 |
| CD44 | 1.00 | [1.00, 1.01] | 0.48 |

## Notes

**Conflict of interest:** The authors declare no potential conflicts of interest.

#### Summary of Updates

Full preprint

## References

1. American Cancer Society. Cancer Facts & Figures 2019, 2019

2. Noone A, Howlader N, Krapcho M, et al: SEER cancer statistics review, 1975–2015. Bethesda, MD: National Cancer Institute, 2018

3. Kohn EC: Invasion and Metastases. Holland-Frei Cancer Medicine 6th edition, 2003

4. Pollard JW: Defining Metastatic Cell Latency. N Engl J Med 375:280–282, 2016

5. Shachar SS, Muss HB: Internet tools to enhance breast cancer care. NPJ Breast Cancer 2:16011, 2016

6. Ravdin PM, Siminoff LA, Davis GJ, et al: Computer Program to Assist in Making Decisions About Adjuvant Therapy for Women With Early Breast Cancer. J Clin Oncol 19:980–991, 2001

7. Mook S, Schmidt MK, Rutgers EJ, et al: Calibration and discriminatory accuracy of prognosis calculation for breast cancer with the online Adjuvant! program: a hospital-based retrospective cohort study. Lancet Oncol 10:1070–1076, 2009

8. Wishart GC, Azzato EM, Greenberg DC, et al: PREDICT: a new UK prognostic model that predicts survival following surgery for invasive breast cancer. Breast Cancer Res 12:R1, 2010

9. Wu X, Ye Y, Barcenas CH, et al: Personalized Prognostic Prediction Models for Breast Cancer Recurrence and Survival Incorporating Multidimensional Data. JNCI J Natl Cancer Inst 109, 2017

10. Kourou K, Exarchos TP, Exarchos KP, et al: Machine learning applications in cancer prognosis and prediction. Comput Struct Biotechnol J 13:8–17, 2015

11. Simon N, Friedman JH, Hastie T, et al: Regularization Paths for Cox’s Proportional Hazards Model via Coordinate Descent. Proc Natl Acad Sci USA 39:1–13, 2011

12. Ishwaran H, Kogalur UB, Blackstone EH, et al: Random survival forests. Ann Appl Stat 2:841–860, 2008

13. Yousefi S, Amrollahi F, Amgad M, et al: Predicting clinical outcomes from large scale cancer genomic profiles with deep survival models. Sci Rep 7:1–11, 2017

14. Kim W, Kim KS, Lee JE, et al: Development of novel breast cancer recurrence prediction model using support vector machine. J Breast Cancer 15:230–238, 2012

15. Delen D, Walker G, Kadam A: Predicting breast cancer survivability: a comparison of three data mining methods. Artif Intell Med 34:113–127, 2005

16. Hanin L, Rose J: Uncovering the natural history of cancer from post-mortem cross-sectional diameters of hepatic metastases. Math Med Biol 33:397–416, 2016

17. Haeno H, Gonen M, Davis MB, et al: Computational Modeling of Pancreatic Cancer Reveals Kinetics of Metastasis Suggesting Optimum Treatment Strategies. Cell 148:362–375, 2012

18. Koscielny S, Tubiana M, Valleron AJ: A simulation model of the natural history of human breast cancer. Br J Cancer 52:515–524, 1985

19. Newton PK, Mason J, Bethel K, et al: A Stochastic Markov Chain Model to Describe Lung Cancer Growth and Metastasis. PLoS One 7:e34637, 2012

20. Retsky MW, Demicheli R, Swartzendruber DE, et al: Computer simulation of a breast cancer metastasis model. Breast Cancer Res Treat 45:193–202, 1997

21. Iwata K, Kawasaki K, Shigesada N: A dynamical model for the growth and size distribution of multiple metastatic tumors. J Theor Biol 203:177–186, 2000

22. Benzekry S, Tracz A, Mastri M, et al: Modeling spontaneous metastasis following surgery: an in vivo-in silico approach. Cancer Res 76:535–547, 2016

23. Koscielny S, Tubiana M, Le MG, et al: Breast cancer: relationship between the size of the primary tumour and the probability of metastatic dissemination. Br J Cancer 49:709–15, 1984

24. Lavielle M: Mixed Effects Models for the Population Approach: Models, Tasks, Methods and Tools. Chapman and Hall/CRC, 2014

25. de Mascarel I, Debled M, Brouste V, et al: Comprehensive prognostic analysis in breast cancer integrating clinical, tumoral, micro-environmental and immunohistochemical criteria. SpringerPlus 2015 4:1 4:528, 2015

26. Gourgou-Bourgade S, Cameron D, Poortmans P, et al: Guidelines for time-to-event end point definitions in breast cancer trials: results of the DATECAN initiative (Definition for the Assessment of Time-to-event Endpoints in CANcer trials). Ann Oncol 26:873–879, 2015

27. Stekhoven DJ, Bühlmann P: MissForest—non-parametric missing value imputation for mixed-type data. Bioinformatics 28:112–118, 2012

28. Ishwaran H, Kogalur UB: Random Survival Forests for R. Rnews 7:25–31, 2007

29. Ishwaran H, Kogalur UB: randomForestSRC: Random Forests for Survival, Regression, and Classification (RF-SRC). 2019

30. Ishwaran H, Kogalur UB, Gorodeski EZ, et al: High-Dimensional Variable Selection for Survival Data. J Am Stat Assoc 105:205–217, 2010

31. Spratt JA, von Fournier D, Spratt JS, et al: Decelerating growth and human breast cancer. Cancer 71:2013–2019, 1993

32. Spratt JS, Meyer JS, Spratt JA: Rates of growth of human solid neoplasms: Part I. J Surg Oncol 60:137–146, 1995

33. Commenges D, Jacqmin-Gadda H: Dynamical Biostatistical Models. Chapman and Hall/CRC, 2015

34. Comets E, Lavenu A, Lavielle M: Parameter Estimation in Nonlinear Mixed Effect Models Using **saemix**, an *R* Implementation of the SAEM Algorithm. J Stat Soft 80, 2017

35. Pedregosa F, Varoquaux G, Gramfort A, et al: Scikit-learn: Machine Learning in Python. Mach Learn PYTHON 6

36. Hossin M, Sulaiman MN: A review on evaluation metrics for data classification evaluations. International Journal of Data Mining & Knowledge Management Process 5:1, 2015

37. Harrell FE, Lee KL, Mark DB: Multivariable prognostic models: issues in developing models, evaluating assumptions and adequacy, and measuring and reducing errors. Stat Med 15:361–387, 1996

38. Bellera CA, MacGrogan G, Debled M, et al: Variables with time-varying effects and the Cox model: Some statistical concepts illustrated with a prognostic factor study in breast cancer. BMC Med Res Methodol 10:20, 2010

39. Cardoso F, van’t Veer LJ, Bogaerts J, et al: 70-Gene Signature as an Aid to Treatment Decisions in Early-Stage Breast Cancer. N Engl J Med 375:717–729, 2016

40. Parker JS, Mullins M, Cheang MCU, et al: Supervised risk predictor of breast cancer based on intrinsic subtypes. J Clin Oncol 27:1160–1167, 2009

41. Dowsett M, Nielsen TO, A’Hern R, et al: Assessment of Ki67 in breast cancer: recommendations from the International Ki67 in Breast Cancer working group. J Natl Cancer Inst 103:1656–1664, 2011

42. Yerushalmi R, Woods R, Ravdin PM, et al: Ki67 in breast cancer: prognostic and predictive potential. The Lancet Oncology 11:174–183, 2010

43. Cheang MCU, Voduc D, Bajdik C, et al: Basal-like breast cancer defined by five biomarkers has superior prognostic value than triple-negative phenotype. Clin Cancer Res 14:1368–1376, 2008

44. Rakha EA, Reis-Filho JS, Ellis IO: Basal-like breast cancer: a critical review. Journal of clinical oncology 26:2568–2581, 2008

45. Lian W, Fu F, Lin Y, et al: The Impact of Young Age for Prognosis by Subtype in Women with Early Breast Cancer. Sci Rep 7:1–8, 2017

46. Al-Hajj M, Wicha MS, Benito-Hernandez A, et al: Prospective identification of tumorigenic breast cancer cells. Proceedings of the National Academy of Sciences 100:3983–3988, 2003

47. Li F, Tiede B, Massagué J, et al: Beyond tumorigenesis: cancer stem cells in metastasis. Cell Research 17:3–14, 2007

48. McFarlane S, Coulter JA, Tibbits P, et al: CD44 increases the efficiency of distant metastasis of breast cancer. Oncotarget 6:11465, 2015

49. Sheridan C, Kishimoto H, Fuchs RK, et al: CD44+/CD24-breast cancer cells exhibit enhanced invasive properties: an early step necessary for metastasis. Breast Cancer Research 8:R59, 2006

50. Perou CM, Sørlie T, Eisen MB, et al: Molecular portraits of human breast tumours. Nature 406:747–752, 2000

51. Benzekry S, André N, Benabdallah A, et al: Modelling the impact of anticancer agents on metastatic spreading. Math Model Nat Phenom 7:306–336, 2012

52. van de Vijver MJ, He YD, van’t Veer LJ, et al: A gene-expression signature as a predictor of survival in breast cancer. N Engl J Med 347:1999–2009, 2002

53. Karrison TG, Ferguson DJ, Meier P: Dormancy of Mammary Carcinoma After Mastectomy. J Natl Cancer Inst 91:80–85, 1999

54. Uhr JW, Pantel K: Controversies in clinical cancer dormancy. Proc Natl Acad Sci USA 108:12396–12400, 2011

